# Spontaneous pain dynamics characterized by stochasticity in awake human LFP with chronic pain

**DOI:** 10.1101/2024.02.22.581655

**Authors:** Jihye Ryu, Jonathan Kao, Ausaf Bari

**Author notes:** Correspondence to Jihye Ryu.

## Abstract

Chronic pain involves persistent fluctuations lasting seconds to minutes, yet there are limited studies on spontaneous pain fluctuations utilizing high-temporal-resolution electrophysiological signals in humans. This study addresses the gap, capturing data during awake deep brain stimulation (DBS) surgery in five chronic pain patients. Patients continuously reported pain levels using the visual analog scale (VAS), and local field potentials (LFP) from key pain-processing structures (ventral parietal medial of the thalamus, VPM; subgenual cingulate cortex, SCC; periaqueductal gray, PVG) were recorded. Our novel AMI analysis revealed that regular spike-like events in the theta/alpha band was associated with higher pain; and regular events in the gamma band was associated with opioid effects. We demonstrate a novel methodology that successfully characterizes spontaneous pain dynamics with human electrophysiological signals, holding potential for advancing closed-loop DBS treatments for chronic pain.

---

Chronic pain is a pervasive condition, impacting approximately 20% of the global population^1^ and contributing to 15-20% of medical appointments^2^. Despite its prevalence, there is a limited understanding of the spontaneous fluctuations in pain experienced by patients with chronic pain. These fluctuations, a key symptom for these patients, involve continuous changes in pain intensity and sensation lasting seconds to minutes, and are observed in conditions like irritable bowel syndrome (IBS)^3^, chronic back pain^4^, postherpetic neuropathy^5^, chronic pelvic pain syndrome^6^, and chronic knee osteoarthritis^7^.

Earlier investigations on pain have aimed at elucidating nociceptive neurons^8,9^ and central pain pathways primarily through animal studies^10^. Human neuroimaging studies have advanced this understanding, revealing that chronic pain is associated with abnormalities in network connectivity^11,12^. These studies have recognized the broader involvement of brain networks, extending beyond sensation to include affect and cognition^13,14^, along with the salience network and default mode network^15^.

Among studies focused on spontaneous pain, many have examined the static resting-state characteristics in chronic pain patients, revealing distinct network and connectivity strength with specific structures ^16,17^. However, these endeavors merely offer a snapshot and fail to capture the ongoing pain fluctuations reported by chronic pain patients in a dynamic manner^18^. Given the intricate link between dynamic brain network processes and pain intensity^19,20^, it is important to understand the brain’s dynamic response to pain on a moment-to-moment basis. A small subset of studies sought to dynamically characterize spontaneous pain fluctuations but were limited by neuroimaging techniques ^5,21,22^, which lack temporal resolution necessary to dynamically capture neural activity. For that reason, examining the dynamic mechanisms behind spontaneous pain would greatly improve our understanding of its emergence and, ultimately, advance treatment options for chronic pain.

To achieve this objective, we captured local field potentials (LFP) from structures engaged in pain processing from five awake patients with chronic pain undergoing awake deep brain stimulation (DBS) implantation surgery. Specifically, we recorded from brain areas implicated in pain processing, conceptualized as pain processing hubs - the ventral posteromedial nucleus (VPM) of the thalamus, the periaqueductal gray region (PVG) ^23,24^, and the subgenual cingulate cortex (SCC) ^25^ (Figure 1A). During awake DBS surgery, patients continuously rated their pain levels by moving a red circle along the visual analog scale (VAS) line displayed on a screen, using a trackpad or buttons for about 10 minutes (Figure 1B). The continuously recorded pain levels were used to segment the data based on varying pain intensity. These segments were analyzed by comparing two consecutive segments and examining the changes in neural activity as pain intensity changed (Figure 1C).

**Figure 1.**
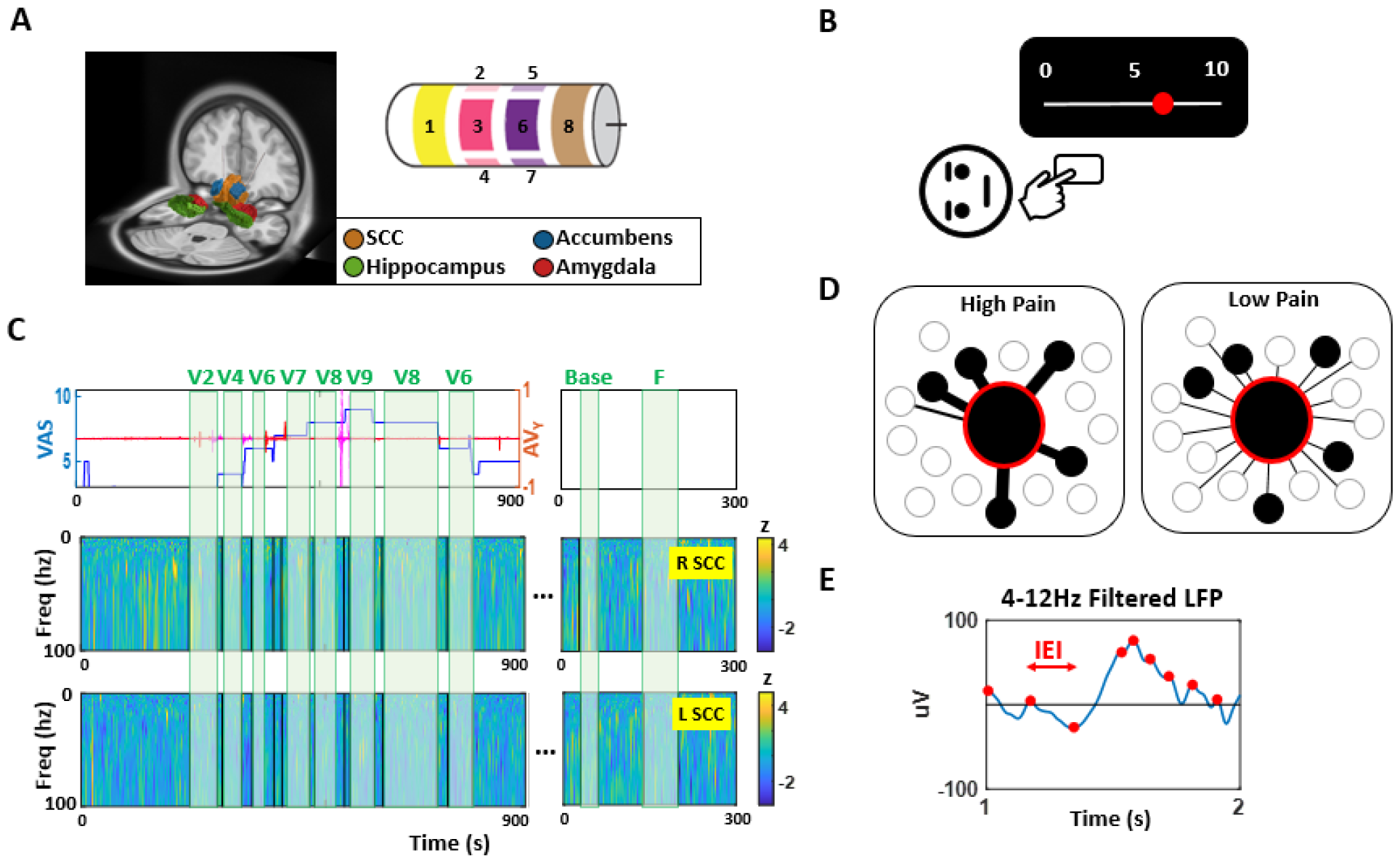
LFP recording, behavioral task, and computational method. **A. (Left)** The DBS lead of a representative patient (P5) is targeting the left and right of the subgenual cingulate cortex (SCC), reconstructed using ^28^. **(Right)** The schematic diagram of the DBS lead depicts a full-ring contact at both the distal end (channel 1) and the proximal end (channel 8) of the cable. In between, there are contacts with equally spaced directional contacts (channel 2-7). **B**. The patient continuously rated their pain level on a VAS from 0 to 10 by moving a red circle using either a trackpad or a button box. **C. (Top)** A representative patient (P5) continuously rated the pain level (blue). Because this patient had low back pain, pain was induced by raising the leg ^29^, and leg raises were recorded with accelerometers attached at the ankle (red). Instances of brief leg raises were excluded from analysis. **(Middle, Bottom)** The spectrograms of the recorded LFP signal were normalized by z-score. The pain VAS trajectory and their corresponding spectrograms for all other patients are shown in Figure S1. **D**. We hypothesize that lower pain levels are characterized by signal irregularity, while higher pain levels exhibit regularity. We conceptualize that irregularity reflects diverse information flow from both pain and non-pain-related sources, while greater regularity indicates focused pain-related information. The schematic diagram illustrates this concept, where black circles indicate pain-related information sources, white circles indicate non-pain-related sources, and line thickness reflects the amount of information flow. **E**. We demonstrate a novel computational method to characterize pain dynamics, which involves extracting the inter-event-intervals (IEI) - time between peaks in the LFP time series - and assessing its stochastic patterns at different pain levels. **Abbreviation:** VAS, visual analog scale; SCC, subgenual cingulate cortex; V pain rating number on a VAS (e.g., V1 denotes pain rating 1 on a VAS)

To characterize pain dynamics, we present a novel computational method in which we quantify the stochasticity in the LFP time series. We conceptualize irregularity to signify diverse (pain- and non-pain-related) information flow from nearby structures, while greater regularity to indicate a more focused flow of pain-related information ^11,26,27^ (Figure 1D). We therefore hypothesize that irregularity in the LFP would characterize a lower level of pain, and regularity to reflect a higher level of pain. Our method involves extracting the time between two consecutive peaks of the theta/alpha bandpass filtered LFP, termed inter-event-interval (IEI) (Figure 1E), which is essentially an instantaneous theta/alpha frequency, as it is an inverse of the corresponding frequency cycle. Here, we quantify the level of regularity by assessing the IEI stochasticity with auto-mutual information (AMI), a nonlinear analogue to auto-correlation. Here, a high AMI would suggest greater regularity, while a low AMI would suggest irregularity. We demonstrate this novel methodology to effectively capture the dynamic changes in response to shifts in pain experience. We further compare the performance of this characterization method with conventional electrophysiology metrics to gain insights into the neural patterns and identify essential features in characterizing spontaneous pain dynamics.

## Materials and Methods

### Participants

Three patients with trigeminal neuropathic pain (participant ID - P2, P3, P4) and two patients with chronic low back pain (participant ID - P1, P5), undergoing a bilateral or unilateral deep brain stimulation (DBS) implantation surgery, participated in this study. The target(s) of the DBS lead included the ventral-parietal-medial (VPM) of the thalamus, periventricular gray (PVG), and/or subgenual cingulate cortex (SCC) (Figure 1A). All subjects provided written informed consent to participate in a research study approved by the institutional review board (IRB) at the University of California, Los Angeles. Details of the patient demographics along with their relevant clinical information can be found in Table 1.

**Table 1.** Participant demographic, relevant clinical information, and change in pain events.

| Participant ID |  | P1 | P2 | P3 | P4 | P5 |
| --- | --- | --- | --- | --- | --- | --- |
| Diagnosis |  | CLBP | TNP | TNP | TNP | CLBP |
| Sex |  | M | F | M | M | M |
| Age |  | 68 | 64 | 67 | 67 | 50 |
| Weight (kg) |  | 79 | 97 | 96 | 103 | 135 |
| Recording Site<sup>1</sup> |  | <u>B SCC</u>,<br>OFC,<br>PFC | <u>R VPM</u> | <u>L VPM, L</u><br><u>PVG</u> | <u>L VPM</u>,<br>OFC, PFC | <u>B SCC</u>,<br>OFC, PFC |
| Bipolar Reference Channels<sup>2</sup> | | 1 -<br>$\mu(2,3,4)$ | $\mu(2,3,4)$-<br>$\mu(5,6,7)$ | VPM:1-<br>$\mu(2,3,4)$<br>PVG:1-$\mu(2,3)$ | $\mu(2,3,4)$-<br>$\mu(5,6,7)$ | $\mu(2,3,4)$-<br>$\mu(5,6,7)$ |
| Sampling Rate (Hz) |  | 5500 | 5500 | 5500 | 4800 | 5500 |
| Condition<sup>3</sup> | Spontaneous (N) | 1 | 1 | 0 | 4 | 7 |
| | $\Delta$ VAS<sup>4</sup> | 1→0 | 9→3 | - | 1→0, 8→0,<br>8→4, 6→4 | 2→1, 3→2,<br>4→3, 6→5,<br>5→4, 7→6,<br>8→7 |
|  | Fentanyl (N) | 0 | 2 | 1 | 0 | 1 |
|  | Bolus (mcg) | n/a | 200<br>(alfentanil<sup>5</sup>),<br>50 | 12.5 | n/a | 25 |
|  | Propofol (N) | 0 | 1 | 1 | 1 | 0 |
|  | Bolus (mg)<br>(infusion rate<br>ug/kg/min) | n/a | 50 mg<br>(125) | 30mg<br>(50) | 30mg<br>(25) | n/a |

<sup>1</sup>Recording sites are listed, but only areas that are underlined are analyzed for the purpose of the study. <sup>2</sup>Channels were bipolar referenced to capture the local activity of the structure. $\mu$ denotes average of the channels listed in parenthesis. Channel numbers correspond to the location along the DBS lead schematically depicted in Figure 1A.
**Abbreviation.** CLBP, chronic low back pain; TNP, trigeminal neuropathic pain; B SCC, bilateral subgenual cingulate cortex; OFC, orbitofrontal cortex; PFC, prefrontal cortex; R VPM, right ventral posteromedial nucleus of the thalamus; L VPM, left ventral posteromedial nucleus of the thalamus; L PVG, left periventricular gray; ΔVAS, change in visual analog scale pain rating.

**Table 2.**
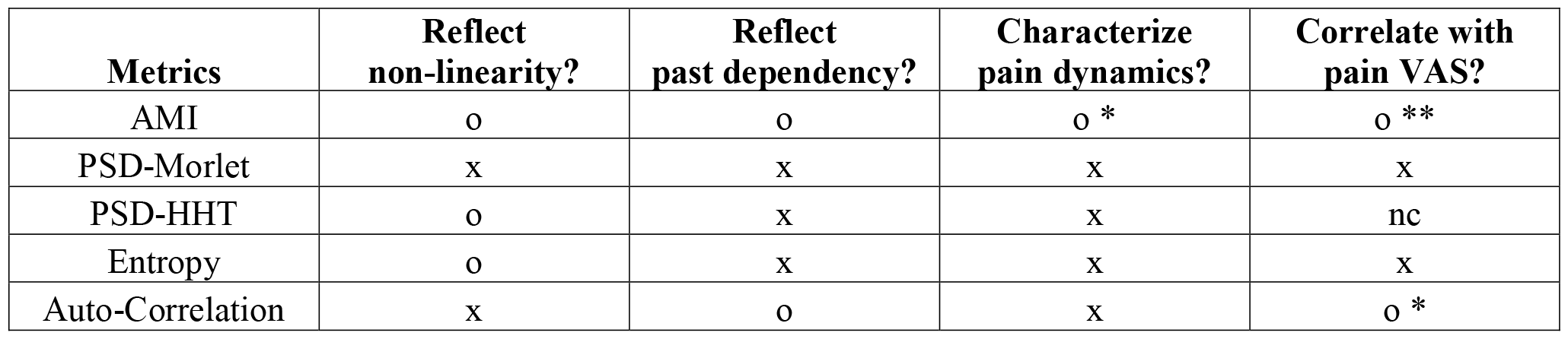
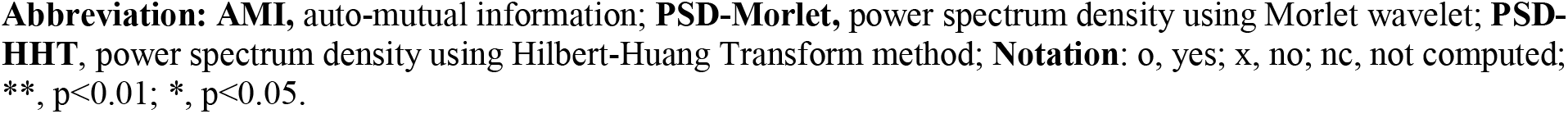
Comparison of metrics for characterizing pain.

| Metrics | Reflect non-linearity? | Reflect past dependency? | Characterize pain dynamics? | Correlate with pain VAS? |
| --- | --- | --- | --- | --- |
| AMI | o | o | o * | o ** |
| PSD-Morlet | x | x | x | x |
| PSD-HHT | o | x | x | nc |
| Entropy | o | x | x | x |
| Auto-Correlation | x | o | x | o * |

**Abbreviation:** AMI, auto-mutual information; **PSD-Morlet**, power spectrum density using Morlet wavelet; **PSD-HHT**, power spectrum density using Hilbert-Huang Transform method; **Notation:** o, yes; x, no; nc, not computed; \*\*, $p < 0.01$ ; \*, $p < 0.05$ .

### Behavioral Task

The patient used a visual analog scale (VAS) displayed on a monitor, along with a trackpad (P1-P4) or button box (P5), to continuously indicate their pain level. Each patient moved a red dot along the VAS scale, which ranged from 0 to 10. The trackpad enabled continuous ratings between 0 and 10, while the button box facilitated discrete integer rating. Patients continuously assessed their pain levels for a duration of 5 to 15 minutes. Afterwards, they were given either fentanyl or propofol based on the clinical protocol reflecting their specific needs. The recording time varied depending on how much the patient’s pain level fluctuated. If a patient had no pain (e.g., patient P3), the pain rating ended after 5 minutes. However, if a patient’s pain rating fluctuated significantly (e.g., patient P4, shown in Figure S1-D), they continued rating their pain for up to 15 minutes. For the two chronic back pain patients (P1, P5), we also utilized intermittent leg raising to induce low back pain similar to the endogenous pain experienced by these patients ^29^. To record the time and extent of the leg raise, we strapped 2 opal inertial measurement unit (IMU) movement sensors (APDM, USA) to the patients’ ankles. The pain rating trajectory of each patient, along with the timing of leg raises for the two chronic low back pain patients are shown in Figure 1A and S1.

### Surgery and data acquisition

During an awake DBS surgery, intraoperative recordings were conducted for research purposes. For all patients, local field potentials (LFP) at the DBS target (P1: bilateral SCC; P2: right VPM; P3: left VPM and PVG; P4: left VPM; P5: bilateral SCC) were recorded from the DBS lead in its final implant position. Patients P1 and P5 had DBS leads manufactured by Abbott (Model 6173ANS, 1.27 mm lead body diameter, inter-contact distance 1.5mm; Abbott Laboratories, Chicago, IL, USA), while the other patients had DBS leads manufactured by Medtronic (Model B3301542 or B3301542M, 1.36 mm lead body diameter, inter-contact distance 2.2mm; Medtronic, Inc., Minneapolis, MN, USA). All DBS leads have a total of 8 electrode contacts and are designed as those shown in Figure 1A. Based on the channel labels in the diagram of Figure 1A, channels 1 and 8 are the most distal and proximal ones respectively, and have full contact. Between them, there are equally spaced 3 directional contacts each at 2 levels, ranging from channels 2 to 4 and channels 5 to 7. LFPs were recorded using these amplifiers: NeuroOmega (AlphaOmega Engineering, Nazareth Israel) for patients P1, P2, P3, P5, Matlab/Simulink software connected to an amplifier (g.Tec, g.USBamp 2.0) for patient P4. The signals obtained from the former amplifier were sampled at 5500Hz, and the latter at 4800Hz. The ground and reference signals were registered using alligator clip electrodes attached to the cannula that was used for the DBS implantation.

For patients with chronic back pain (P1, P5), their legs were intermittently raised to simulate their low back pain. We used two Opal IMU movement sensors (APDM, USA) strapped on their ankles to capture the kinematics, specifically the angular velocity, and measure the timing and extent of the leg raise. To synchronize between the electrophysiological and kinematic signals, we utilized an external synchronization equipment (“sync box” provided by APDM, USA). This equipment sent a digital output trigger to the electrophysiological signal amplifier, precisely indicating the start and end times of the IMU sensor recording.

For patients who received either fentanyl or propofol, we manually recorded the timing of the medication administration. Due to potential human error, the precise timing of the administration may have a deviation less than 5 seconds. However, since both medications typically take around less than a minute to take effect, such minor imprecision is not expected to significantly impact the overall results presented in this study.

## Analysis

### Preprocessing

LFPs recorded from each of the 8 channels of the DBS lead were applied with a 60 Hz notch filter (band-stop IIR, filter order 80) and then visually examined for artifacts and excess noise. To reduce noise from the 8 recorded channels, we performed the following data processing steps. First, we averaged the signals from the 2nd level (channels 2-4) and 3rd level (channels 5-7) of the DBS lead (Figure 1A). Then, we applied bipolar referencing between adjacent levels as follows: Channel 1 was bipolar referenced against the average of channels 2-4; the average of channels 2-4 was bipolar referenced against the average of channels 5-7; and the average of channels 5-7 was bipolar referenced against channel 8. Given that the distal part of the DBS lead (channel 1) is typically implanted closest to the targeted structure, we prioritized examining the bipolar referenced signal of channel 1 against the average of channels 2-4. However, in certain cases where channel 1 exhibited excess noise (i.e., lacking 1/f structure in the power spectrum), we resorted to examining the bipolar referenced signal of the average of channels 2-4 against the average of channels 5-7. Additionally, for patient P3, the PVG LFPs recorded showed excess noise in channels 2-5. As a result, we opted to bipolar reference the signal between channel 1 and the average of channels 6-7. The exact channel numbers used for the bipolar referencing are provided in Table 1. Once the channels were identified for further analysis, we conducted visual inspections to exclude segments with electrical artifacts. These artifacts were characterized by a sudden and distinct change in amplitude (>500uV), lasting for more than 1 second. We removed such segments, and the total length of removed segments did not exceed more than 5 seconds in duration for any of the patients.

To segment the data by epochs based on the patient’s pain levels, we considered both the pain rating and the verbal reports conveyed during the recording. As an example, for patient P1, who expressed feeling minimal pain throughout the recording, we excluded data associated with leg raises to avoid any interference from the effects of leg raise, and only selected times when the patient verbally expressed change in pain. We also considered the acclimation time for patients who used the trackpad or a button box to indicate their pain rating. Patients using the trackpad took around 1-2 minutes to get acclimated, while patients using the button box took less than 1 minute. For all patients, we excluded the initial acclimation period from the analysis. Furthermore, we addressed the issue of excessive fluctuations in VAS ratings registered through the trackpad (due to patients’ unintentional motions) by applying a moving average method with a smoothing factor 0.25 (“smoothdata.m” function in Matlab). This allowed us to visually identify time segments with different pain levels. Lastly, for patient P5, whose leg was raised intermittently to induce pain, we excluded times when the leg was in motion. Specifically, we selected time segments when there were no leg movements (i.e., angular speed of the angle was less than 0.01 degrees/second) for at least 10 seconds. After carefully considering these criteria, which are 1) ensuring that the pain ratings aligned with the patient’s verbal reports, 2) accounting for their familiarity with the trackpad or button box, 3) identifying stable VAS ratings, and 4) excluding data with leg movements, we proceeded to segment the data into epochs of different pain levels obtained from this process. To capture the impact of pain medication, we created an epoch by extracting data starting from 1 minute after the administration of the medication and spanning approximately 45 to 60 seconds. The segmentation is visually depicted by green boxes in Figure 1C for patient P4 and in Figure S1 for all other patients. At the end, these resulted in 33 epochs across all patients (median epoch duration 42 seconds; IQR = 17, 60 seconds).

### Auto mutual information (AMI) based on inter-event-interval (IEI)

After dividing the dataset into epochs, we proceeded to calculate the auto mutual information (AMI) for each epoch and observed the changes in AMI when pain was lower. AMI is considered a nonlinear analog of the auto-correlation function, and employs information theory ^30^to quantify the predictability of future points in a time series based on past points. The relationship between past and predicted future time points is reflected in the diminishing AMI as the time delay increases, providing insights into the system’s complexity ^31^. Notably, research has demonstrated that higher AMI values (indicative of reduced complexity) are linked to with various conditions such as schizophrenia^32^, idiopathic dilated cardiomyopathy ^33^, Alzheimer’s disease ^34,35^, multiple organ dysfunction syndrome and heart failure ^36^. In this context, we computed the epoch-specific AMI to characterize the dynamics of pain, and to gain insight into the obtained results in relation to complexity.

The formula to compute AMI is identical to mutual information (MI), a measure that quantifies the probabilistic dependency between signals from two different sources. In the context of AMI, the dependency is computed within a single source, where one signal is temporally shifted by a specified time delay in relation to the other signal. To elaborate, let’s consider X to represent the IEI series from the first time window (i.e., future points) and Y to represent the IEI series from the time window with a time delay (i.e., past points). AMI, denoted as *I*_*XY*_ in the formula below, quantifies the extent to which the uncertainty of X can be reduced by having knowledge of Y. This concept is defined in equation (1), wherein *H* represents an entropy function that quantifies the degree of uncertainty.

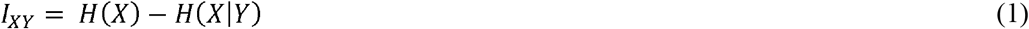

Here, *H*(*X*) is the entropy of X, which signifies the inherent uncertainty within X, as defined in equation (2). The term *H*(*X*|*Y*) represents the conditional entropy of X given Y, depicting the remaining uncertainty in X when Y is known, as defined in equations (3-5).

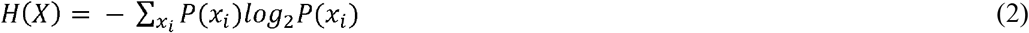

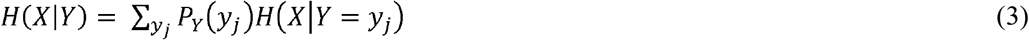

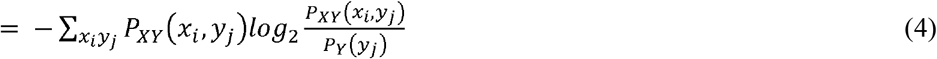

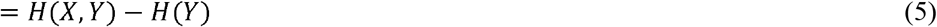

Based on equations (2) and (5), I_XY_ can be simplified as equation (6), which was used to calculate the AMI values in this study.

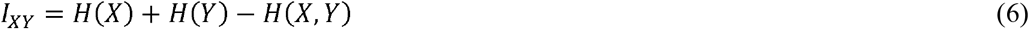

To do this, first, the LFP was initially down-sampled to 400Hz. Subsequently, we computed two distinct sets of AMIs, each based on different bandwidths. Note, we chose to initially down-sample the data to 400hz, because the minimum IEI was found to be 2.5ms based on the raw data that was filtered at 30-80hz (gamma), and the duration of a single frame within a 400hz data equals 2.5ms.

The first AMI was computed using data that underwent bandpass filtering within the theta and alpha frequency range (4-12Hz; FIR, 200th order zero-phase, transition width 0.2). These filtered time series were then partitioned into 5-second intervals, which were paired with a 5-second window delayed by 175ms (Figure 2A). For each of these time windows, local peaks were identified (marked as red dots in Figure 2B). The duration between consecutive local peaks (derived with “findpeaks.m” function in Matlab; minimum peak prominence 0, threshold 0, minimum peak distance 0), referred to as inter-event-intervals (IEIs), was extracted for every time window. Given the sequence of IEIs, these were represented as a time series aligned with the 400hz frame size (Figure 2C). For each paired time windows, probability distributions were constructed for the IEIs of the preceding time window (referred to as P(IEI) in Figure 2D or P(X) in the equations above) and for the IEIs of the subsequent time window (referred to as P(IEI’) in Figure 2D or P(Y) in the equations above). Additionally, a joint probability distribution (P(IEI,IEI’) in Figure 2D or P(X,Y) in the equations above) was constructed based on the IEI and IEI’ occurrences. These distributions enabled us to compute the components *H*(*X*), *H*(*Y*), *H*(*X, Y*), which ultimately produced a single AMI value. Paired time windows were shifted forward with 50% overlap across the entire epoch, resulting in a series of AMI values (rightmost graph in Figure 2D). The mean AMI was selected for comparison between epochs. Note, the variables *i* and *J* in the equations (2-4) correspond to the bins encompassing the range of IEIs, spanning from 10 to 130 frames with increments of 10 frames (i.e., 25ms to 325ms in 25ms increments). We chose the time window length of 5-second to gather around 50 IEI datapoints, enabling a meaningful frequency distribution, while capturing the dynamical changes in AMI over the epoch duration (about 45 seconds). We chose the shift size of 70 frames (175ms), as it effectively represents the IEI’s mean duration, and optimally differentiates the epochs with differing pain intensity states (See Figure S2 for different lengths of shift size).

**Figure 2.**
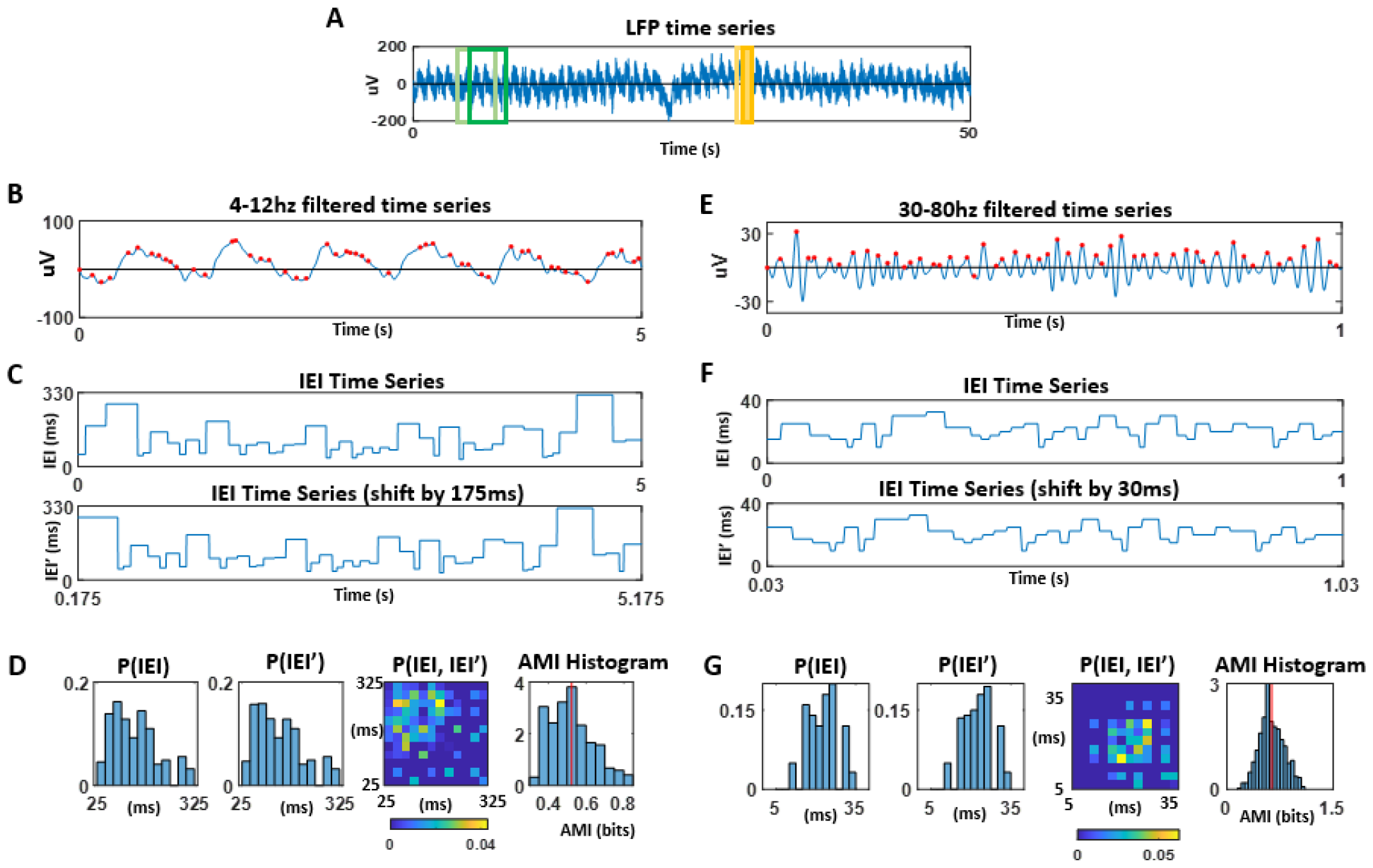
Auto mutual information (AMI) computation pipeline. **A**. LFP from each recorded structure was divided into epochs. Within each epoch, two types of windows were chosen for analysis. The first window had a duration of 5 seconds, and it was paired with another window delayed by 175ms (shown in green). These windows were used for theta/alpha bandwidth analysis (4-12hz). Another type of window had a duration of 1 second and was paired with another window delayed by 30ms (shown in orange). These windows were used for gamma bandwidth analysis (30-80hz). For both 5-second and 1-second paired windows, these were slid forward with 50% overlap across the entire epoch time frame. **B**. To analyze within the theta/alpha bandwidth, the 5-second time windows were bandpass filtered at 4-12hz. The local peaks (denoted in red) within these windows were marked to compute the inter-event intervals (IEI), which is defined as the time between peaks. **C**. IEI time series were created for the paired time windows. **D**. Within the 5-second time window, probability distributions of the inter-event intervals (IEIs) were computed for the paired window, along with the joint probability distribution of paired-IEIs. This process yielded a single AMI at each slid time point, and the AMI values across the entire epoch were collected. The mean of the AMI (shown in red line) was extracted for later comparison between epochs. **E**. To analyze within the gamma bandwidth, 1-second time windows were bandpass filtered at 30-80hz, and local peaks were extracted (shown in red dots). **F**. Similar to C., time series data for IEI were generated for the corresponding windows. **G**. Probability distributions and a joint probability distribution of the IEIs were computed and used to calculate the AMI at each time point. The AMIs from all time points were collected, and the mean of the AMIs (shown as a red line) was extracted for later comparisons.

The second AMI was computed using data that was bandpass filtered within the gamma frequency range (30-80Hz; FIR, 200th order zero-phase, transition width 0.2). These filtered time series were then partitioned into 1-second intervals, which were paired with a 1-second windows delayed by 30ms. Then, local peaks were identified (marked as red dots in Figure 2E) to compute the IEIs for every time window. Given the sequence of IEIs, these were represented as a time series (Figure 2F), and the paired time window’s IEI time series were used to construct the necessary probability distributions to compute the AMI at a given time point (Figure 2G). Paired time windows, overlapping by 50%, were shifted across the entire epoch. Mean AMI within each epoch was then compared to another (rightmost graph in Figure 2G). Note, the distribution bins (variables *i* and *J* in the equations (2-4)) spanned from 5 to 14 frames with increments of 1 frame (i.e., 12.5ms to 35ms in 2.5ms increments). We chose the time window length of 1-second to gather around 50 IEI datapoints, enabling a meaningful frequency distribution, while capturing the dynamical changes in AMI over the epoch duration. We chose the shift size of 12 frames (30ms), as it effectively represents the IEI’s mean duration, and optimally differentiates the epochs with differing pain intensity states (See Figure S2 for different shift size).

We highlight that while the AMI metric has found utility in prior neural studies (e.g.,^33–37^), variations exist in defining the constituent variables (i.e., X and Y in equations 1-6) for constructing probability distributions. Indeed, converting a continuous signal like electrophysiological data into an appropriate discrete probability space is a challenging task. To that end, we introduce an innovative approach rooted in kinematics studies ^38,39^ to define constituent variables. Diverging from conventional AMI computations involving amplitude binning, our method concentrates solely on the timing information, disregarding amplitude details. By reducing the raw time series to a binary spike train, this approach effectively simplifies the computation while presenting a versatile solution suited to diverse signal modalities. Notably, prior studies (e.g., electrocorticography^40^, EEG^41^, voice^42^, magnetometer^43^) have successfully harnessed the inter-event interval (IEI) information to characterize a range of biophysical signals, showing its adaptability across different modalities.

### Statistical comparisons of AMI between low and high pain levels

To assess the AMI changes when pain is lower, we analyzed the data in two ways, by: 1) combining all lower pain instances across all brain structures (VPM, PVG, SCC) and 2) separating the lower pain instances by structure. When these data were separated by structure, we are left with a small sample size. For that reason, when we combined the structures, we performed the Wilcoxen signed rank test for zero median, and when we separated the data by structures, we performed a series of permutation tests to see whether there were significant changes in the signal’s AMI.

### Wilcoxon signed rank test for zero median

Using each instance of lower pain, we conducted a Wilcoxon signed rank test for zero median (non-parametric analog of a one-sample t-test) to determine if the difference between matched samples comes from a distribution with a median of zero. To do this, we collected the mean AMI values from each lower pain epoch and its adjacent (subsequent or prior) higher pain epoch (from here on, we will refer these as the *paired epochs*). Then, we subtracted the lower pain epoch’s mean AMI value from the higher pain epoch’s mean AMI value for each paired epochs (low pain AMI – high pain AMI). This non-parametric test identifies statistical significance when the paired epochs exhibit an AMI difference departing from zero. Note, based on a power analysis considering our available data points, achieving significant results requires a total sample size of 16 for a two-sided one-sample t-test with a power of 0.8 and an alpha of 0.05. While the test applied to AMI comparisons during spontaneous lower pain is adequate for reaching statistical significance (N=21), the sample size is insufficient for AMI comparisons during lower pain induced by fentanyl (N=8) and propofol (N=4). We complement these findings with permutation tests performed separately for each structure below.

### Permutation test

To address the challenge posed by a relatively small sample size when analyzed per structure, we performed a two-sided permutation test involving 1000 permutations. The permutation test aimed to assess whether the mean difference between the lower and higher pain states within each paired epochs significantly differed from zero. For the theta/alpha range AMI comparisons, we randomly selected 3 AMI values without replacement from each epoch. Note, within a single epoch, there were 115 AMI values on average (ranging from 19 to 409). We then shuffled the AMI values between the epoch pairs (higher and lower), and computed the difference between mean AMI from the shuffled-lower and shuffled-higher epochs. We repeated this 1000 times, to generate a null distribution of 5000 datapoints (5 paired epoch x 1000 repetition) for the spontaneous lower pain in VPM, 4000 (4 paired epoch x 1000 repetitions) for fentanyl-induced in VPM, 3000 (3 paired epoch x 1000 repetitions) for propofol-induced in VPM, 16000 (16 paired epoch x 1000 repetition) for spontaneous lower pain in SCC, 4000 (4 paired epoch x 1000 repetition) for fentanyl-induced in SCC, and 1000 (1 paired epoch x 1000 repetition) for propofol-induced in PVG. The observed mean difference between the paired epochs were then compared to this null distribution, allowing us to determine statistical significance. For the gamma range AMI comparisons, we repeated the above permutation test methods, but by randomly selecting 25 AMI values without replacement from each epoch. Within a single epoch for gamma band-passed signal, there were 125 AMI values on average (ranging from 27 to 125).

### Kruskal-Wallis nonparametric one-way analysis of variance (ANOVA)

To confirm that the AMI values do not differ between different types of lower pain, we performed the Kruskal-Wallis test on the AMI values across spontaneous, fentanyl-induced, and propofol-induced conditions at both higher and lower pain states. Given the skewed non-normal distribution of the AMI, and unequal size and variance between conditions, we chose this non-parametric test that allowed to address these data features.

### Comparison metrics

To illustrate the utility of AMI and assess its critical features in characterizing pain dynamics, we also applied established metrics commonly used to analyze neural activity. These metrics capture different signal attributes such as non-stationarity, non-linearity, and dynamic past dependency. By comparing the efficacy of pain characterization between AMI and these metrics, we identified the critical attributes to capture the neurophysiological patterns of pain.

### Power Spectrum

Using the preprocessed LFP, we conducted power spectrum analysis utilizing the BOSC algorithm^44^. This examination covered individual frequencies ranging from 1 to 100Hz with a step width of 1Hz and employed 6th order Morlet wavelets. To standardize the power time series, we applied z-scoring to each frequency by normalizing across the entire recording. We then extracted the z-scores from epochs and compared these across different epochs. By analyzing the changes in z-scores derived from the Morlet wavelet power spectrum, we aimed to determine whether a metric capable of mitigating the stationarity assumption proves effective in characterizing pain dynamics.

As an alternative to the BOSC method, we also computed the power spectrum using the Hilbert-Huang Transform (HHT)^45^ with frequency limits from 1 to 100hz. Intrinsic mode functions (IMF), that are necessary to compute the HHT power spectrum, were set with default parameters specified in the Matlab functions emd.m and hht.m. After obtaining the Hilbert spectrum for the entire recording, we applied z-scoring to each frequency for normalization and compared the z-scores corresponding to different epochs. Using the HHT method, we aimed to determine whether a metric that relaxes both linearity and stationarity constraints is effective in characterizing pain dynamics.

To statistically compare across frequency bands ranging from 1 to 100 Hz, we applied the aforementioned permutation test at each individual frequency level. Additionally, we accounted for multiple comparisons by employing the Bonferroni correction. This involved dividing the significance level (alpha 0.05) by the total number of frequency bins (100) to maintain an adjusted threshold for statistical significance. This enabled us to assess whether a non-dynamic metric with relaxed assumptions of stationarity and/or linearity could effectively characterize pain dynamics.

### Auto-Correlation

Using the bandpass-filtered and epoch-segmented LFP, we computed auto-correlation for each paired epochs, adhering to the identical time window lengths and sliding percentage as specified in the AMI approach. We performed the Wilcoxen signed rank test based on the mean auto-correlation values from the paired epochs, as was done with AMI analyses. Using this metric, we aimed to evaluate whether a linear and dynamic metric that captures past dependency such as auto-correlation exhibited a positive correlation with the pain VAS.

### Entropy

Using the spike trains generated for AMI calculation, we extracted the inter-event interval (IEI) data points within each epoch. We then computed entropy using the same binning parameters as in the AMI method. We computed the entropy from paired epochs and compared the change in entropy using the Wilcoxen signed rank test. Notably, this approach captured the static “uncertainty” observed throughout the epoch and did not capture dynamic past dependency. The objective of this comparison was to determine whether a static information-theory metric, such as entropy, captures pain more effectively than the dynamic AMI metric.

### Statistical correlation with pain VAS

To determine if there’s a correlation between the pain VAS and the aforementioned metrics, we calculated the Kendall rank correlation coefficient (referred to as Kendell’s τ coefficient) between pain VAS and each metric. Kendall’s coefficient is a suitable choice since it’s a non-parametric test commonly used to gauge ordinal associations.

In addition, we assessed whether the direction of change in pain (increasing vs. decreasing in time) makes a difference in the AMI. To do this, in contrast to how we assessed lower pain by comparing any two sequential epochs, we separated by those that are increasing pain in time versus those that are decreasing in time. After separating the AMI values by increasing pain and decreasing pain, we conducted a Kendall’s correlation between the corresponding pain ratings and the AMI for each set.

## Results

### Lower pain is characterized by irregularity (reduced AMI) in the theta/alpha frequency band

The AMI derived from the LFP within the theta/alpha frequency bands did not differ across the three conditions – spontaneous vs. fentanyl-induced vs. propofol-induced (during high pain, χ^2^(2,30)=3.1, p=0.21; during low pain, χ^2^(2,30)=3.6, p=0.16). However, when we compared between the paired epochs, AMI was significantly less than zero when pain was lower (Wilcoxon signed-rank test for zero median). This AMI reduction was observed both when pain was lower spontaneously (n=21; p=0.04) and when fentanyl was administered (n=8; p=0.02) but not under propofol administration (n=4; ns) (Figure 3A). The individual data points illustrating the mean AMI for the paired epochs separated by recording site per patient and by conditions are shown in Figure 3B. A majority of datapoints were below the unity line, indicating that AMI is generally lower when pain is lower.

**Figure 3.**
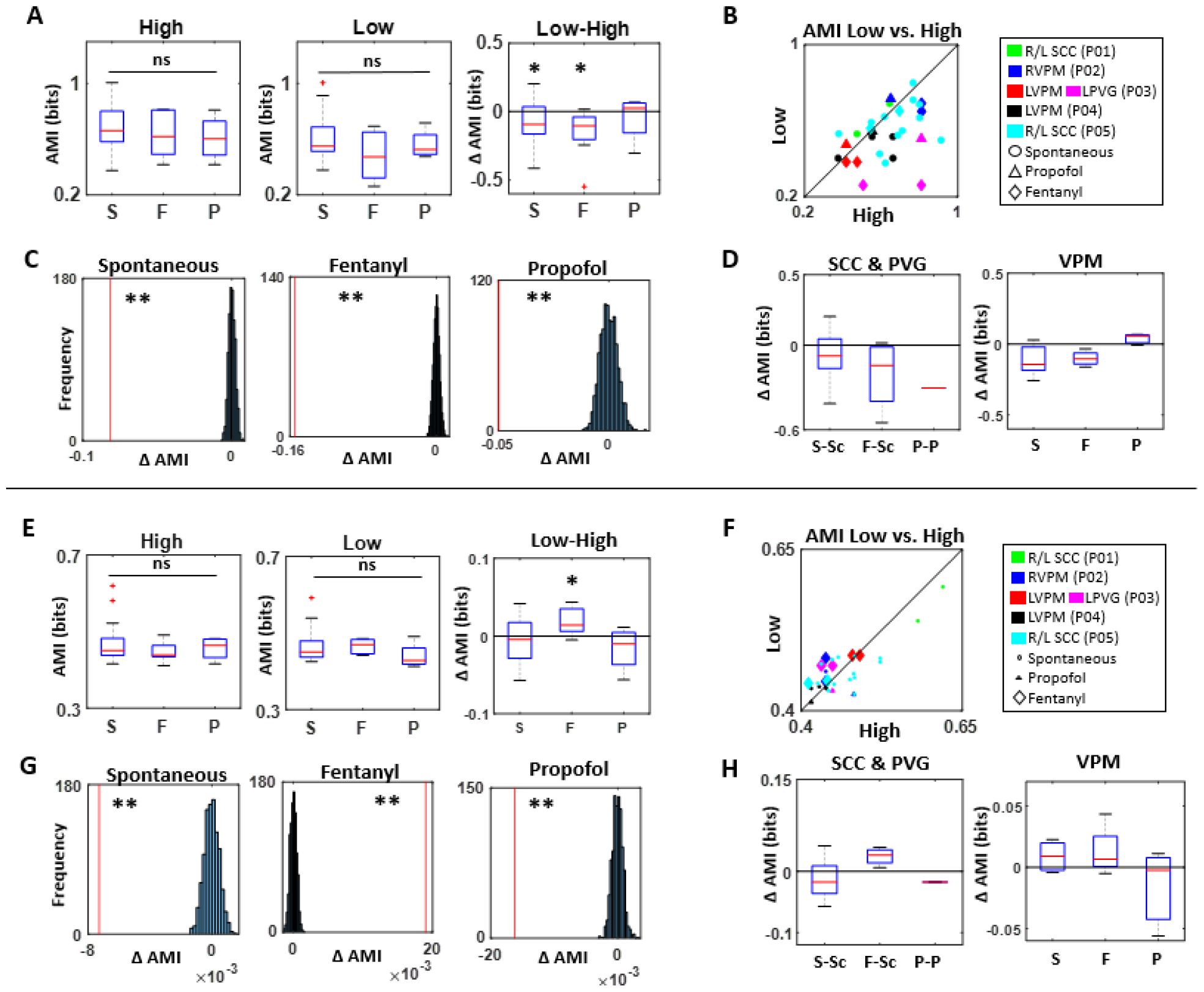
Results of Theta/Alpha Frequency Band (A-D) and Gamma Band (E-H) Dynamics. <u>Theta/Alpha Band Dynamics:</u> **A**. AMI values did not show differences between S, F, and P conditions under high (left) and low (middle) pain states, based on Kruskal-Wallis non-parametric ANOVA test. When changes in AMI are examined for each lower pain instances (i.e., difference in paired epochs) using the Wilcoxon signed-rank test for a zero median (right), AMI is reduced under S and F conditions, but not under P condition. **B**. Mean AMI values are plotted for each paired epochs - low and high pain states - per patient (differentiated by marker color) and by conditions (differentiated by marker shape). Each data point corresponds to a single paired epoch. **C**. The change in AMI from each condition is assessed using permutation tests, showing significant reduction for each. The red line denotes the observed AMI change, while the distribution represents the shuffled null distribution. **D**. When the results of AMI change are assessed separately by structures, the decrease in AMI is consistent for S and F conditions across all structures. <u>Gamma Band Dynamics:</u> **E**. AMI values do not show differences between S, F, and P conditions under high (left) and low (middle) pain states, based on the Kruskal-Wallis non-parametric ANOA test. However, when changes are examined per paired epochs using the Wilcoxon signed-rank test for a zero median (right), AMI values increase only under F condition. **F**. Similar to **B**, single data points are plotted for each paired epochs differentiated by marker color (patient) and shape (condition), where F conditions are denoted by a larger diamond shape. **G**. Similar to **C**, permutation test results are performed for each condition, revealing an increase in AMI under F condition, and a reduction in AMI under S and P conditions. **H**. Similar to **D**, the Wilcoxon signed-rank test is performed separately by structure, revealing an increase in AMI values under F conditions for all structures. **Abbreviation**: **S**, spontaneous lower pain condition; **F**, fentanyl based lower pain condition; **P** propofol administered condition; **S-Sc**, spontaneous lower pain condition in SCC; **F-Sc**, fentanyl-based lower pain condition in SCC; **P-P**, propofol-induced condition in PVG; **L-H**, Low-High; Δ**AMI**, change in AMI from lower pain (i.e., AMI from low pain – AMI from high pain); **\*\***, significance p<0.01; **\***, significance p<0.05; **ns**, non-significant.

To address the limitation posed by a small sample size, permutation testing was also performed on these AMI data points. These tests revealed a consistent AMI reduction when pain was lower across all three conditions (p<0.01) (Figure 3C). Given that these data points encompass the VPM (n=5 for S; n=4 for F; n=3 for P), SCC (n=16 for S; n=4 for F; n=1 for P), and PVG (n=1 for P) structures, we also segregated these structures, and confirmed that the directional shift in AMI remains uniform across structures, with the exception of the case involving propofol administration in the VPM (Figure 3D).

It is worth noting that we also conducted tests for the temporal shift between time windows at various lengths (ranging from 25ms to 275ms). The results indicate that a temporal shift of 175ms (as shown in the main results) provides the optimal distinction between high and low pain states (Figure S2 A). Furthermore, we explored different frequency bands, including theta only, alpha only, alpha/low beta, and theta/alpha/low beta, and found that the theta/alpha bands offer the most effective characterization of pain dynamics (Figure S2 C).

### Effect of opioid pain medication is associated with regularity (increased AMI) in the gamma frequency band

To examine the effect of opioid pain medications on LFPs within the VPM, PVG and SCC, we performed a similar analysis as the above, but limited the bandwidth to the gamma frequency range. Here, the AMIs did not exhibit any differences between the three conditions, whether it was during high pain (χ^2^(2,30)=2.19, p=0.3) or low pain (χ^2^(2,30)=0.76, p=0.7) states. However, we observed an increase in AMI upon fentanyl-induced lower pain, (Wilcoxon signed-rank test for a zero median) (n=8; p=0.02), and no change in spontaneous (n=21; p=0.24) and propofol-induced (n=4; p=0.38) conditions (see Figure 3E). Figure 3F presents individual data points depicting the mean AMI for each paired epochs denoted by recording site, per patient, and condition.

We also conducted permutation testing on the change in AMI datapoints per paired epochs. These tests consistently revealed an increase in AMI under fentanyl-induced lower pain (p<0.01), and AMI reduction for the other two conditions (p<0.01), as illustrated in Figure 3G. Similar to the above analysis, we also segregated these by recorded structures and confirmed that the directional shift in AMI under fentanyl-induced lower pain remained the same for VPM (n=4) and SCC (n=4) (see Figure 3H).

We also conducted tests for temporal shifts between time windows of various lengths, ranging from 15ms to 35ms, and found that 30ms temporal shift (as shown in the main results) provided the best distinction between high and low pain states (see Figure S2 B). Furthermore, we explored whether restricting the bandwidth to low gamma (30-60hz) or high gamma (60-80hz) would be more effective in characterizing pain dynamics, but found the 30-80hz range to be the most optimal (see Figure S2 D).

### Dynamical past dependency captures pain dynamics

AMI based on the IEI parameter is a novel method that captures dynamical past dependency and nonlinearity of a single LFP signal, and we demonstrated how this successfully characterized pain dynamics. Next, we aimed to understand which features of this method contributed to an effective characterization of pain dynamics. To explore this further, we applied a variety of conventional metrics using datasets obtained from the spontaneous condition (as these included pain VAS scores) and aimed to identify key features to capture the pain dynamics.

First, we found that AMI doesn’t merely characterize pain dynamics (i.e., AMI reduction corresponds to lower pain) in categorical manner (as shown in Figure 3A), but that it is correlated with the pain VAS (τ=0.50; p<0.01). We further validated this for both rising (τ=0.64; p<0.01) and falling pain trends (τ=0.79; p<0.01) and confirmed that AMI mirrors the overall pain intensity level (Figure 4A). We also compared this per patient P4 and P5 – the two patients who experienced a wide range of pain VAS during the recording – and confirmed that their AMI values positively correlated with their pain VAS at a marginal (p<0.1) level on the left side of the recorded structure for P4 and P5 (P4: τ = 0.80, p=0.08; P5: τ = 0.69, p=0.02). P5’s right side structure did not show significant positive correlation (τ = 0.50, p=0.14) (Figure 4B).

**Figure 4.**
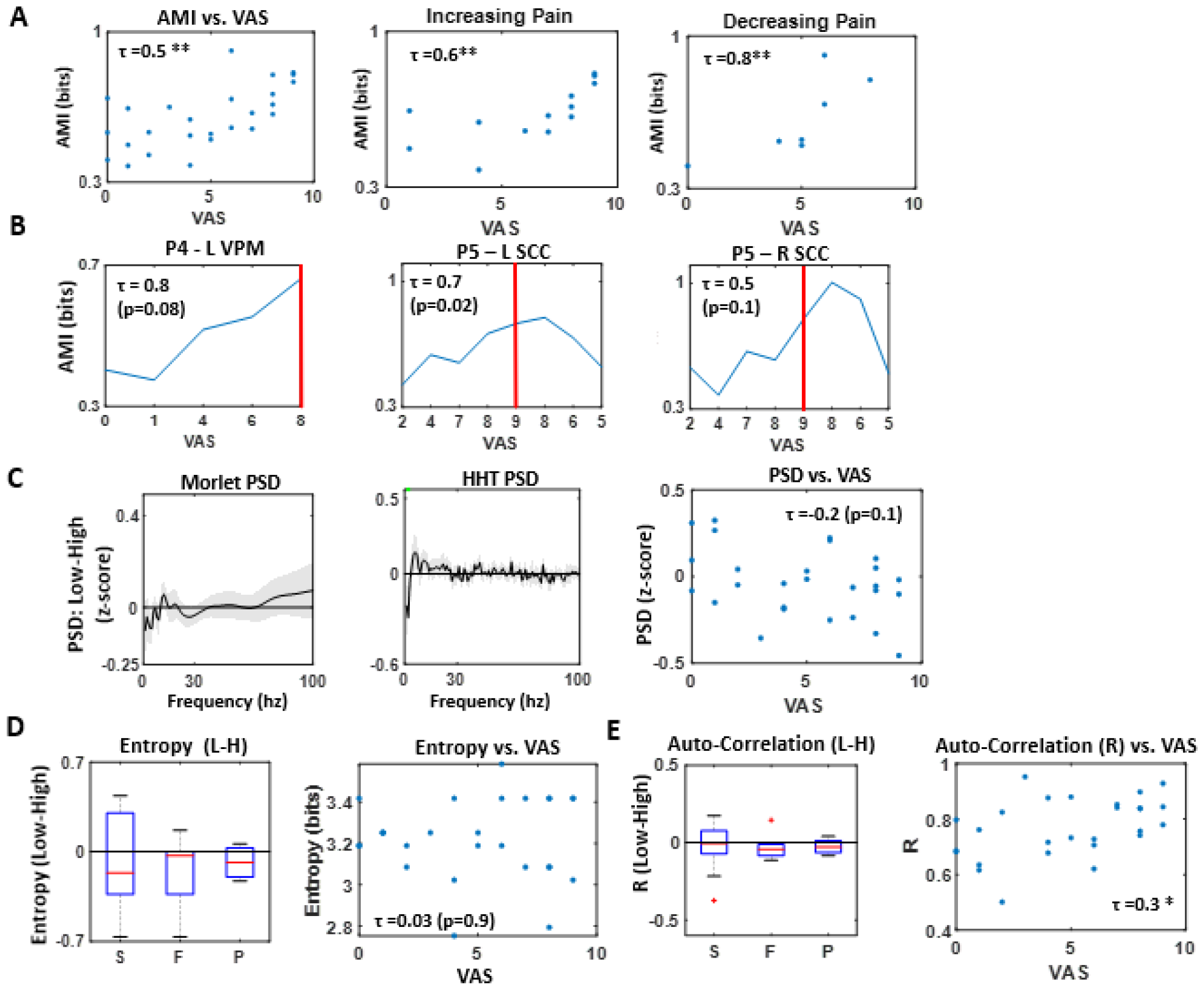
Comparison of metrics in characterizing pain dynamics. **A**. AMI positively correlated with pain VAS (left), and was indifferent to whether the pain was increasing (middle) or decreasing (right) in time. **B**. AMI positively correlated against pain VAS individually for patient P4 (left) and P5 (middle, right), and left structures showed a stronger correlation than the right structure. For ease of reading, for P4 the x-axis (pain VAS) is ordered by pain scores; whereas for P5, the x-axis follows a temporal order. The red line marks the highest pain score. **C**. Changes in PSD (z-score) using the Morlet wavelet (left) and the HHT method (right) following spontaneous lower pain are plotted across all frequency bands. The shaded area represents the standard error of the mean. In PSD-HHT at 2 Hz, significance was observed at p < 0.05 without multiple corrections. No significant correlation was found between PSD and pain VAS (right). **D**. Changes in entropy following spontaneous lower pain were not evident (left), nor did they exhibit any correlation with the pain VAS (right). **E**. Changes in auto-correlation following spontaneous lower pain were not observed across all conditions, although the auto-correlation value exhibited a positive correlation with pain VAS scores. **Abbreviation**: **L VPM**, left ventral-parietal-medial of the thalamus; **L SCC**, left subgenual cingulate cortex; **R SCC**, right subgenual cingulate cortex; **PSD**, power spectrum density; **HHT**, Hilbert-Huang Transform; **R**, auto-correlation coefficient; τ, Kendall’s tau correlation coefficient; **L-H**, Low-High; **S**, spontaneous lower pain condition; **F**, fentanyl based lower pain condition; **P** propofol administered condition; **\*\***, significance p<0.01; **\***, significance p<0.05; **ns**, non-significant.

To compare with other metrics, we also examined whether power spectrum density (PSD) could effectively capture pain dynamics. This assessment was carried out using both the Morlet wavelet and the Hilbert-Huang Transform (HHT) methods. It is noteworthy that while neither of these methods reflects past dependency, the latter does capture nonlinearity. Our findings revealed that neither of the PSD methods demonstrated effective characterization of pain dynamics. Furthermore, we conducted a correlation between the normalized (z-scored) Morlet wavelet PSD values and the pain VAS scores but did not find any significant patterns (τ = -0.23, p=0.11) (Figure 4C). This suggests that the inclusion of dynamic past dependency may be a critical factor in characterizing pain dynamics.

To assess the importance of nonlinearity, we compared entropy values between paired epochs, as the entropy metric captures the nonlinearity feature but does not account for past dependency. Here, our findings showed that entropy did not effectively capture pain dynamics across conditions (S, p=0.57; F, p=0.25; P, p=0.50), nor did it correlate with the pain VAS (τ = 0.03, p=0.9), This indicates that nonlinearity is not a critical feature (see Figure 4D) to characterize pain dynamics. To investigate the importance of past dependency, we compared the auto-correlation values between paired epochs. This metric does not reflect nonlinearity features but does capture past dependency within a specific time window (in this case, 5 seconds). As a result, we did not observe auto-correlation to effectively capture pain dynamics under all conditions (S, p=0.86; F, p=0.20; P, p=0.38). However, we did find a positive correlation between auto-correlation values and the pain VAS (τ = 0.33, p=0.02) (Figure 4E). This finding suggests that past dependency may play a critical role in characterizing pain dynamics, although the presence of linearity might diminish its effectiveness when compared to the current AMI method.

## Discussion

Our goal was to apply a novel method - AMI based on IEIs – on the LFP signals at the VPM, PVG, and SCC to characterize pain dynamics. As we conceptualized AMI as a measure of regularity in the signal (indicated by increased past dependency), we discovered a link between pain and regularity (high AMI) within the theta/alpha band. Additionally, we found that opioid-based pain medication led to regularity (high AMI) within the gamma band. This discovery marks the first to *dynamically* characterize spontaneously occurring pain fluctuations with human electrophysiological signals.

In addition, our novel computational approach has several merits that potentially make it a valuable pain biomarker for application in closed-loop DBS treatment. First, we demonstrate the sufficiency of using signals from a single site to characterize pain. This was possible because we recorded from a neural hub that processes diverse pain-related information ^23–25^. Notably, we could not achieve this when calculating AMI in the prefrontal cortical sites (shown in Figure S2 D). Therefore, our findings differ from viewpoints favoring the use of signals from multiple sites as an effective way to characterize pain ^49^. Additionally, the AMI method we introduced is computationally more efficient than typical machine-learning methods ^46,50^, which require extensive data and training (lasting hours to days). Note, we merely needed about 10 seconds worth of LFP time series to compute the AMI. Our method also surpasses the efficiency of prior neural studies using the AMI metric (e.g., ^34,35,51^), which typically depend on signal amplitude values. Here, we reduce the LFP time series to binary spike trains for computing the inter-event intervals (IEIs) thus simplifying the computation. Considering these strengths, we believe this novel methodology holds significant potential to serve as a robust biomarker for the development of closed-loop DBS for chronic pain.

We further explored which features of this AMI method most contributed to its effective characterization in pain dynamics. To do this, we employed various metrics that capture different features of the signal and assessed their effectiveness in characterizing pain dynamics. As a result, we found the auto-correlation metric to be the most suitable alternative for characterizing pain dynamics. As auto-correlation measures the extent of past dependency in the signal, we consider that a decrease in past dependency (i.e., irregularity) in the signal plays a critical role in effectively characterizing lower pain. Moreover, considering our current AMI method’s superior performance compared to auto-correlation, our speculation is that its effectiveness stems from its nonparametric and nonlinear approach in quantifying past dependency.

We interpret these results with the conceptualization that irregularity in the LFP signal represents diverse (pain- and non-pain-related) information flow from nearby structures, whereas greater regularity reflects a more focused flow of pain-related information. Specifically, we consider that in the absence of pain, the signal becomes more irregular and less reliant on past data (i.e., lower AMI), as diverse information about the external world is constantly communicated, updating the signal moment-to-moment. Conversely, during the experience of pain, analogous to how an individual in pain concentrates on the painful sensation, the communication primarily involves pain-related information originating from within the internal body. In this case, the signal tends to hold onto its internal pain information, impeding the input about the external world or other non-pain-related factors. With this viewpoint, we hypothesized that higher pain would be associated with more signal regularity and lower pain with signal irregularity, and we found this to be the case within the theta/alpha frequency band. Our results align with prior research, where lower variability was found in fMRI BOLD signals from somatosensory cortex and PVG during pain states ^20^, and in EEG signals among sensory-over responsive individuals ^41^. Interestingly, we find an inverse trend between the spectral power and AMI, as we note an increase in theta/alpha power coincided with lower AMI (irregularity) upon pain reduction, and a decrease in gamma band power (Figure S3 A, B) coincided with higher AMI (regularity) following fentanyl administration. However, we also note that these two metrics (power spectrum and AMI) do not exhibit a significant negative correlation when directly compared against each other (Figure S3 C). From this, we conjecture that when new information flows into a site within a specific oscillatory bandwidth, the signal becomes more irregular (lower AMI), while more oscillatory power emerges within that frequency band. To gain a deeper understanding of this hypothesis, investigating the neural connectivity between the pain processing hub structures and neighboring sites would provide a more comprehensive view of this concept of information flow.

We also interpret our findings on past dependency within the context of complexity. Although infrequent, AMI has been applied in electrophysiological studies as a measure of complexity, representing rich interconnectedness and information content ^31^. Specifically, higher AMI would indicate lower complexity, and has been linked to various diseases (e.g., Alzheimer’s disease ^34,35^, schizophrenia ^32^, idiopathic dilated cardiomyopathy ^33^, multiple organ dysfunction syndrome^36^). Regarding our results, we interpret that higher AMI (found in higher pain instances) is associated with a less complex brain state, indicating sparsely interconnected information flow within the theta/alpha band. Interestingly, we observed that pain relief induced by opioid medication increases AMI within the gamma band, implying reduced complexity. This suggests that opioids may introduce an artificial information flow leading to less complex interactions within the gamma band. While drawing definitive conclusions is premature, we argue that our findings provide insights into the frequency bandwidths worth exploring in future studies involving opioid pain medication, to deepen our understanding of the complexity of brain dynamics in such contexts.

We recognize that the study’s small sample size, consisting of 5 participants, limits the generalizability of our findings. However, it’s crucial to underscore that each individual data point originates from an extensive dataset (ranging approximately from 500 to 2500 datapoints) acquired from each patient. Consequently, these individual data points possess notable statistical power and serve as meaningful summary statistics on their own. Furthermore, we acknowledge the variability in epoch lengths, both among patients and within paired epochs, which results in an uneven sample size from which the mean AMIs are derived. This approach was unavoidable as we aimed to maintain an observational approach within a naturalistic context. Still, we assert that the AMI values, rooted in the mean of a stochastic process with more than 100 datapoints, would not be too sensitive to the sample size (compared to statistical parameters like variance).

In future studies, it would also be valuable to conduct this research within a more controlled setting, especially when investigating the impact of pain medication. Additionally, the calculation of AMI in our study involved selecting parameters such as frequency bandwidth, bin size, and time window overlap, which would affect the resulting AMI values. We used a data-driven approach to derive these selections, going through multiple trial-and-error iterations (depicted in Figure S3 and S4 B) to discover optimal parameters that effectively differentiated the pain states. For subsequent research, we suggest applying these selected parameters to a larger group of participants and confirm whether these parameter choices hold true for a wider range of individuals. We also recognize that the patients in this study monitored pain in an intraoperative setting, which is different from a more naturalistic everyday setting. To validate our findings, assessing LFP in a naturalistic environment, especially in patients with externally recorded DBS implants, would enhance the authenticity of pain experiences in chronic pain patients. Lastly, gaining a deeper understanding of the underlying neural activity could be achieved by simultaneously recording single-unit spike activity alongside population-based activity. This approach could eventually lead to the development of a model-based understanding of this phenomenon.

In summary, our study presents a novel AMI computational method for characterizing pain dynamics through the analysis of electrophysiological signals in the VPM, SCC, and PVG regions. This discovery offers significant potential for advancing the development of closed-loop DBS treatments for individuals with chronic pain, ultimately leading to enhanced care and improved outcomes for a wide range of chronic pain patients.

## Supplemental Figures

**Figure S1.**
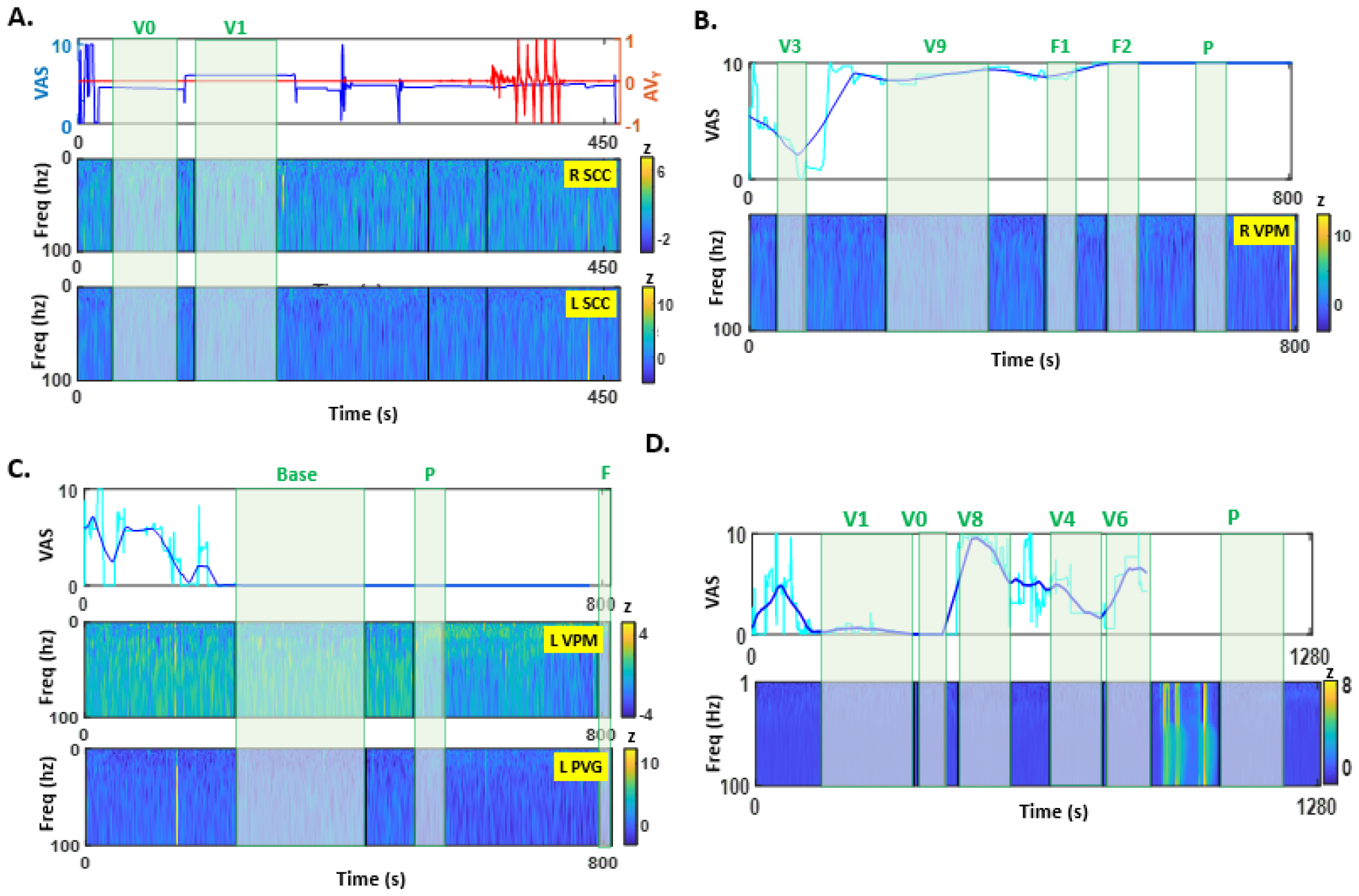
Pain VAS alongside the corresponding spectrogram and epoch specifications for each patient. **A. Patient P1**. (top) Pain VAS is denoted by blue, while leg angular velocity along the Y-axis is represented by red. Initially, despite the VAS being marked at 4, we labeled it as “V0” because the patient verbally reported no pain. A subsequent increment of +1 in pain VAS was noted as “V1”. **B. Patient P2**. The patient’s trackpad trajectory is depicted in cyan, and the smoothed VAS is shown in blue. **C. Patient P3**. This patient verbally indicated the absence of pain and was accordingly administered propofol, followed by fentanyl as per clinical protocol. **D. Patient P5**. The patient’s incremental fluctuations in pain are shown in blue and angular velocity along the Y-axis are shown in red. Subsequently the patient performed an unrelated experiment for approximately 15 and then was administered with fentanyl due to experienced pain. **Abbreviation. V**, pain VAS; **F**, fentanyl administration; **P**, propofol administration; **R SCC**, right subgenual cingulate cortex; **L SCC**, left subgenual cingulate cortex; **R VPM**, right ventral parietal medial of the thalamus; **L VPM**, left ventral parietal medial of the thalamus; **L PVG**, left periventricular gray.

**Figure S2.**
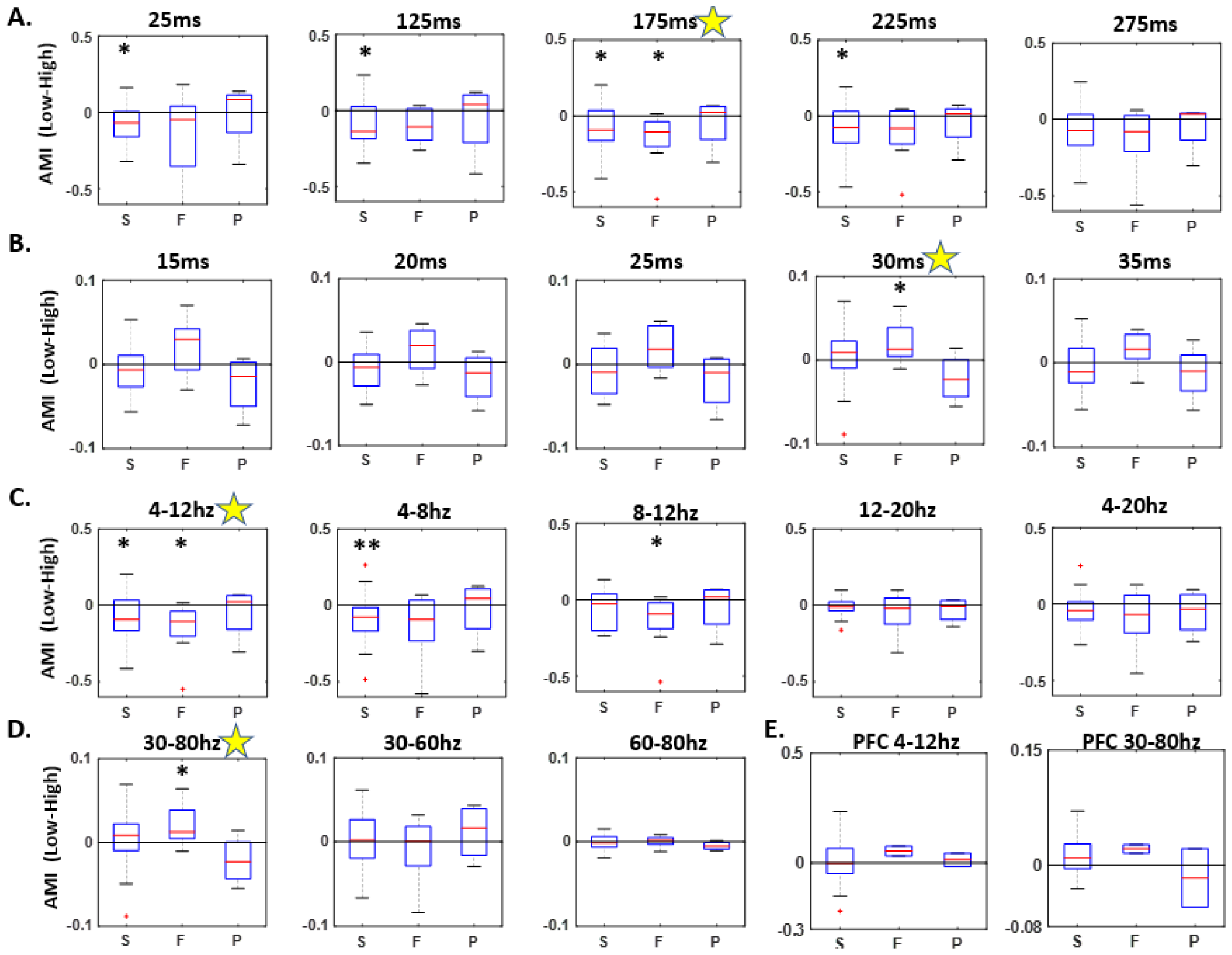
Change in AMI under varying parameters. **A**. When applying a theta/alpha bandpass filter to the LFP, we computed the change in AMI resulting from pain relief across different time-delayed windows (ranging from 25ms to 275ms). It was determined that the 175ms time delay yielded the most optimal characterization of pain relief. **B**. For the LFP filtered within the gamma frequency band, the change in AMI stemming from fentanyl intake was best characterized using a time delay of 30ms. **C**. The computation of change in AMI from pain relief involved exploring different frequency bandwidths within the lower frequency range. Notably, the theta/alpha band emerged as the most suitable for characterizing pain relief from both spontaneous and fentanyl-induced instances. **D**. Similarly, change in AMI from pain relief was calculated across different frequency bandwidths within the higher frequency range. Results indicated that the 30-80Hz band provided the most effective characterization of pain relief from fentanyl. **E**. We extended the AMI comparison to the prefrontal cortex. No significant change in AMI was detected for either the theta/alpha or gamma bands upon experiencing pain relief.

**Figure S3.**
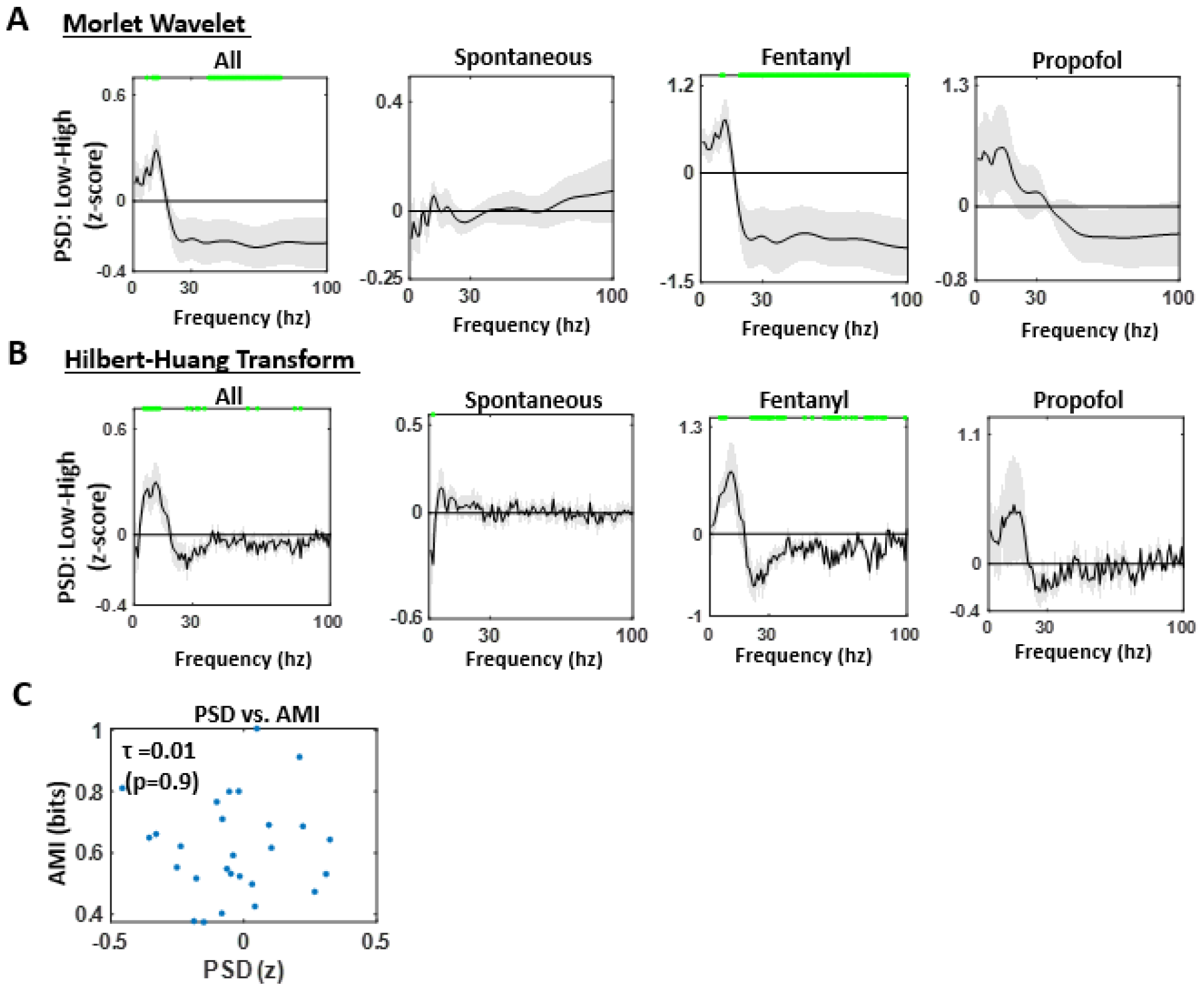
Change in power spectrum from pain relief and its relation to AMI. **A**. The change in power spectrum (z-score) from pain relief is depicted using Morlet wavelets for all types of pain relief, and categorized into spontaneous, fentanyl-induced, and propofol-induced. The shaded region represents the standard error of the mean. Instances of statistical significance at p<0.05, without corrections for multiple comparisons, are indicated in green. However, no statistical significance was observed across all frequency ranges when corrections were applied. **B**. The change in power spectrum (z-score) from pain relief is presented using the Hilbert Huang Transform method. This method exhibits a comparable pattern to the Morlet wavelet approach, showing an observable increase in power spectrum within the lower frequency range (around the theta band) and a decrease in the gamma band range. Despite this consistency, no statistical significance was detected upon applying corrections for multiple comparisons. **C**. When comparing the power spectrum density (PSD) and AMI values, no direct correlation is evident between the two metrics.

**Figure S4.**
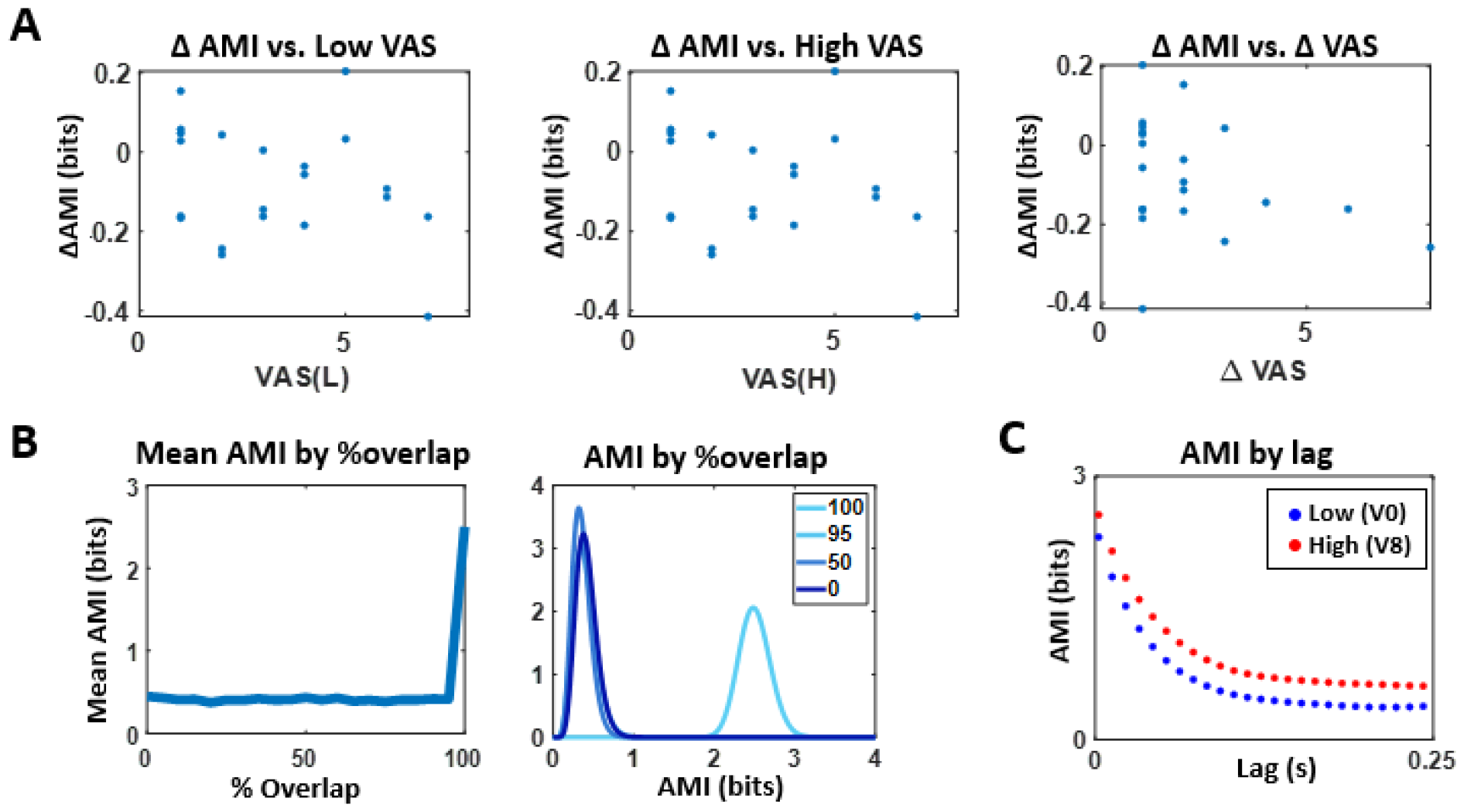
Miscellaneous analyses on AMI computation. The magnitude of change in AMI does not exhibit correlation with VAS measurements. Specifically, the change in AMI resulting from pain relief does not correlate with the lower pain VAS, as evidenced by the relationship between ΔAMI and the corresponding lower pain VAS (τ = -0.20, p=0.24), nor does it correlate with higher pain VAS (τ = -0.20, p=0.24). Furthermore, the change in AMI does not show correlation with the magnitude of change in pain VAS, as indicated by the correlation between ΔAMI and ΔVAS (τ = -0.24, p=0.18). **B**. Between two time windows with time delay, observations revealed that when the two windows experienced no time delay (indicated by 100% overlap), the mean AMI was the highest. Nonetheless, when the overlap reduced, alterations in time delay did not result in significant changes to overall AMI values. Thus, while we identified the optimal time delay for AMI computation in this study, we believe that variations in time delay would not substantially impact the overall outcomes. **C**. In nonlinear dynamical systems, a steeper decline in the rate of AMI with time delay serves as a normalized complexity measure ^31^. In our present study, we observed an exponential decrease in the AMI function within the lag range of up to 250ms. Consequently, a steeper decline would correspond to lower AMI values.

